# Quantitative three-dimensional nondestructive imaging of whole anaerobic ammonium-oxidizing bacteria

**DOI:** 10.1101/709188

**Authors:** M-W. Peng, Y. Guan, J-H. Liu, L. Chen, H. Wang, Z-Z. Xie, H-Y. Li, Y-P. Chen, P. Liu, P. Yan, J-S. Guo, G. Liu, Y. Shen, F. Fang

## Abstract

Anaerobic ammonium-oxidizing (anammox) bacteria play a key role in the global nitrogen cycle and the treatment of nitrogenous wastewater. These functions are closely related to the unique biophysical structure of anammox bacteria. However, the research on the biophysical ultrastructure of intact anammox bacteria is lacking. In this study, *in*-s*itu* three-dimensional nondestructive ultrastructure imaging of whole anammox cell was performed using synchrotron soft X-ray nano-computed tomography and the total variation-based simultaneous algebraic reconstruction technique (TV-SART). Statistical and quantitative analyses of the ultrastructures of intact anammox bacteria were performed. The linear absorption coefficient values of the ultrastructures of anammox bacteria were calculated and the asymmetric structure of the anammox bacteria was quantified. On this basis, the shape adaptation of the anammox bacteria responses to Fe^2+^ were explored, and the underlying regulation mechanism of Fe^2+^ on anammox bacteria was explored. Furthermore, a promising method to study the biophysical properties of cells in different environments and engineering processes was proposed.

**Graphical Abstract:** 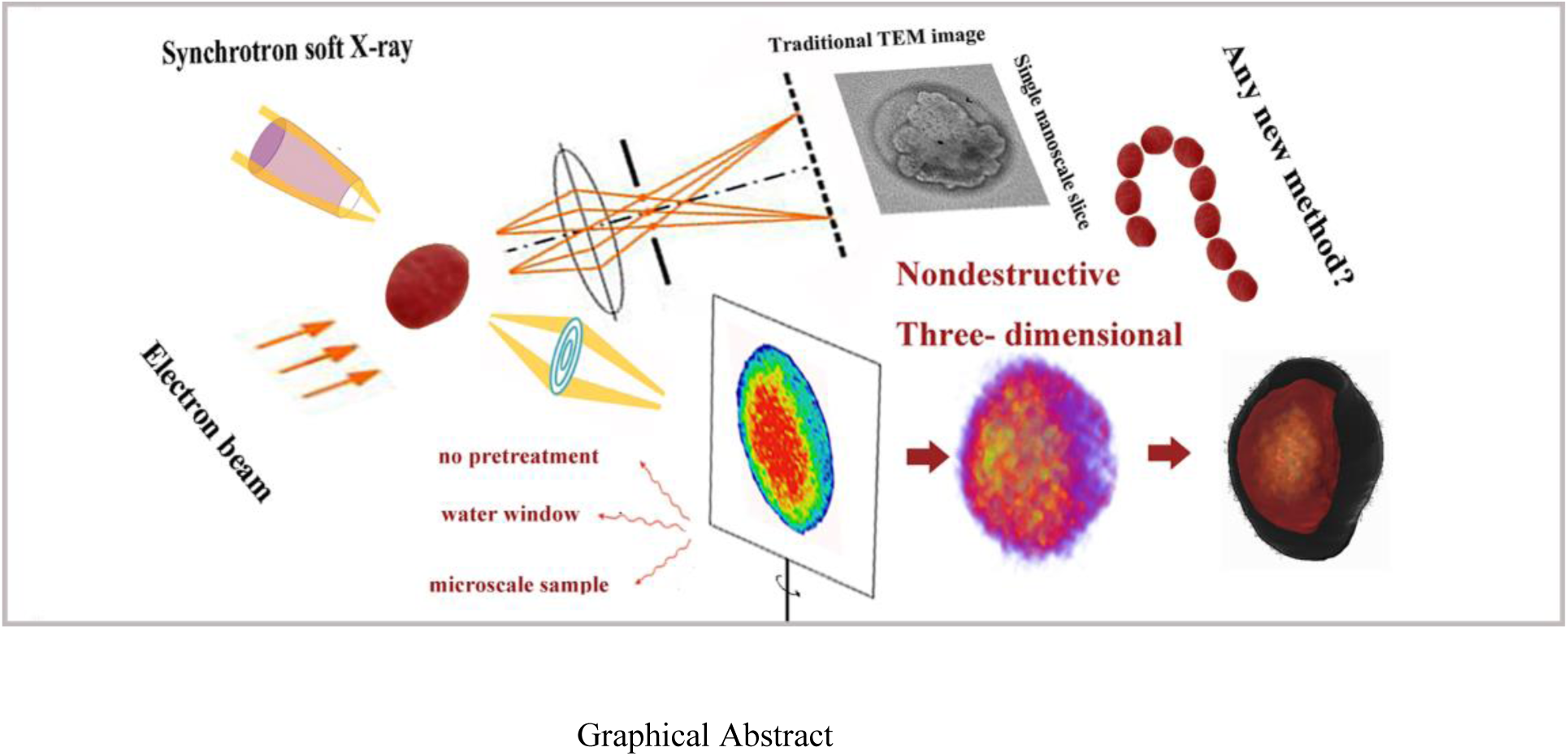

**Statement of Significance:** Anaerobic ammonium-oxidizing (anammox) bacteria play key role in global nitrogen cycle, and this physiological function depends on the unique morphology of anammox bacteria. In this study, synchrotron soft-X ray imaging technique coupled with simultaneous algebraic reconstruction technique with total variation (SART-TV) algorithm were performed to quantify the three-dimensional ultrastructure of the whole anammox bacteria for the first time. On this basis, the shape adaptation and mechanism of the anammox bacteria responses to Fe2+ were explored and a promising method for detecting the physiological properties of anammox bacteria was proposed.

## Introduction

Anaerobic ammonium oxidizing (anammox) bacteria are widespread bacteria in rivers (1), marine sediment (2), paddy soil (3), oil fields (4), and wastewater treatment plant (5). Anammox bacteria are estimated to account for up to 50% of the nitrogen removal from the oceans (6). Moreover, the anaerobic, autotrophic anammox bacteria obtain energy from the oxidation of ammonium with nitrite to nitrogen gas (2, 7, 8). This property makes anammox bacteria suitable for the removal of nitrogen compounds from nitrogenous wastewater due to their environmental friendliness and cost-effectiveness (9). Further, anammox bacteria are prokaryotic organisms but contain a fairly large organelle structure. Anammox bacteria have an irregular, membrane-bounded intracellular compartment called the anammoxosome (10). The anammoxosome organelle is unique and only exists in anammox bacteria (11). The key catabolic metabolism occurs in anammoxosome (11, 12). In addition, anammox membranes contain unique membrane lipids called ladderanes (13). Ladderane lipids are calculated to have a density of 1.5 kg/dm^3^ and are unique to anammox bacteria (14). The dense membrane reduces the leakage of protons and valuable intermediates of the anammox process (15). This complex biological structure makes anammox bacteria unique in terms of biological morphology.

The morphology of microbes is closely related to their function. The multicellular filamentation of *Herpetosiphon* was presumed to be associated with predatory behavior, predation defense and motility in aqueous or soil environments (16). *Caulobacter crescentus* adapts to different environmental challenges by adjusting their shape (17). The stalk of *Caulobacter* was elongated up to 30 *μ*m when faced with phosphate starvation (18) and this stalk elongation structure was hypothesized to enhance phosphate absorption (19). Moreover, *Shewanella alga BrY* adapts to starvation *by* reducing the cell volume from 0.48 to 0.2 *μ*m^3^; the starved cell can be resuscitated to the prior volume in a suitable environment (20). Similar morphological changes were reported in Mn-oxidizing bacteria, i.e., *Halomonas meridiana and Marinobacter algicola* in response to Mn (II)-induced stress. The cell length of *Halomonas meridiana and Marinobacter algicola* increased significantly to improve the Mn (II) oxidizing ability (21). The cell morphology of bacteria can be adapted dynamically depending on environmental conditions, a requirement for optimal survival and growth (17). Thus, morphology research is of great significance. However, the morphology of intact anammox bacteria has not been investigated to date and the shape adaptations of anammox bacteria under different environment conditions remain unclear.

Currently, the most effective and advanced imaging method for anammox bacteria is the transmission electron microscope (TEM) technique. In spite of the successful application of the TEM method for the analysis of the anammox structure, the limited transmission depth, the inherently low contrast, and the destructive pre-treatment including dehydration, embedding, and the ultrathin section (22) made it impossible to date to image the 3D structure of intact anammox bacteria. The development of cryo-electron tomography has resulted in many breakthroughs in the imaging field (23) but the method requires that the sample has to be cut into nanoscale slices (at most up to 500nm). Limited by the weak penetration, cryo-electron tomography is incapable of high throughput imaging for micron-thick cells (24). To date, the structure of the whole anammox bacteria cell has not been determined. The lack of effective *in-situ* imaging methods of whole anammox bacteria limits the exploration of the unique metabolic mechanism of anammox bacteria. Thus, an *in-situ* nondestructive method for the morphological study of intact anammox bacteria is urgently required to reveal the microstructure and related metabolic functions of anammox bacteria.

Synchrotron soft X-ray nano-computed tomography (CT) is an *in-situ* nondestructive imaging technology. It provides a method to obtain a non-destructive 3D image of whole hydrous cells. Synchrotron soft X-ray photons can penetrate hydrated specimens up to 15 *μ*m. This property permits the imaging of the entire intact anammox cell in its natural state without any staining, sectioning, dehydration, or embedding pretreatments (25, 26). The synchrotron soft X-ray wavelength is adjustable and by using the “water window” range from 2.3 nm to 4.4 nm, the absorption of organics is almost an order of magnitude higher than that of the water, thus the organics can be distinguished from the water by natural contrast at 30-nm resolution (26). In addition, this imaging method allows for obtaining the transmission image along the projection direction, as well as providing information on the interior structure of the specimen (26, 27). Thus, unlike other imaging methods, synchrotron soft X-ray nano-CT provides a unique opportunity for imaging intact anammox cells in the *in-situ* state. The synchrotron soft X-ray imaging technique was successfully applied to the 3D mapping of eukaryotic cells, such as the 3D reconstruction of saccharomyces organelles (26), imaging of the structure of hepatitis C virus (HCV) replicon-harboring cells (28), and research of superparamagnetic iron oxide nanoparticle reactions with a breast cancer cell (29). However, to date, the synchrotron soft X-ray nano-CT technique has not been applied to anammox cells; therefore, this topic requires investigation.

In this study, the synchrotron soft X-ray nano-CT imaging technique was used to image an intact anammox bacteria cell. A reconstruction based on the traditional filtered-back projection (FBP) calculation method was used to reconstruct the numerous image slices. The total variation-based simultaneous algebraic reconstruction technique (TV-SART) was employed in this study to optimize the imaging process and obtain the image information. The linear absorption coefficient (LAC) was determined based on the TV-SART algorithm. The 3D ultrastructure of the anammoxosome volume ratio, the eccentricity, and the nanoparticles of an intact anammox cell in its natural hydrous state were explored and the shape adaptations of anammox bacteria in response to environmental changes were investigated.

## Materials and Methods

### Sample preparation for analysis

#### Cultivation and purification of the anammox granules

The anammox bacteria were cultivated in a 2-L expanded granular sludge blanket (EGSB) reactor (30). The reactor has been running steadily for five years and the mineral medium (per liter demineralized water) consist of NH_4_Cl, 764 mg (200 mg/l NH_4_-N); NaNO_2_, 985 mg (200 mg/l NO_2_-N); KHCO_3_, 2.133 g; KH_2_PO_4_, 25 mg; MgSO_4_.7H_2_O, 200 mg; CaCl_2_.2H_2_O, 200 mg,; FeSO_4_, 0.03mM ; and 1 mL of trace element solution I and II as described by Van de Graaf *et al* (31). The hydraulic retention time of the EGSB reactor was 3.7 h and 13 L/d of the mineral medium was degraded by the anammox bacteria in the EGSB reactor. The anammox bacteria extracted from the EGSB reactor were purified by Percoll density centrifugation (32).

#### DNA extraction and PCR amplification from for the granular sludge

Microbial DNA was extracted from the ammonia oxide granular sludge in the EGSB reactor using the E.Z.N.A. ® soil DNA Kit (Omega Bio-Tek). The final DNA concentrations were determined using an ultraviolet-visible (UV-VIS) spectrophotometer (PerkinElmer Lambda 950) and the DNA molecular mass was determined by gel electrophoresis with 1% agarose. The V3-V4 regions of the anammox 16S rRNA gene were amplified with the primer 338F (5’-ACTCCTACGGGAGGCAGCAG-3’) and the primer 806R (5’-GGACTACHVGGGTWTCTAAT-3’) using polymerase chain reaction (PCR) (ABI GeneAmp 9700). The PCR program was as follows: 3 min of denaturation at 95 °C, 27 cycles of 30 s at 95 °C, 30 s for annealing at 55 °C, 45 s for elongation at 72 °C, and a final extension at 72 °C for 10 min. A mixture of 4 μl of 5 × FastPfu Buffer, 2 μl of 2.5 mM dNTPs, 0.8 μl of 5 *μ*M 338F primer and 0.8 μl of 5*μ*M 806R primer, 0.4 μl of FastPfu Polymerase, 0.2μl BSA, and 10 ng template DNA was added to the reaction system; deionized water was added to obtain 20 μl. Two parallel PCR reactions were performed at the same time. The resulting PCR products were extracted from 2% agarose gel and further purified using the AxyPrep DNA Gel Extraction Kit (Axygen Biosciences) and were quantified by QuantiFluor™-ST (Promega).

### Characteristic analysis of the anammox bacteria

#### Diversity analysis of the anaerobic granular sludge

The Illumina MiSeq PE300 platform (Illumina, San Diego, USA) was used for sequencing of the purified amplicons and the raw reads were deposited into the National Center for Biotechnology Information (NCBI) Sequence Read Archive (SRA) database. The operational taxonomic units (OTUs) were clustered with a 97% similarity cutoff using UPARSE (version 7.1 http://drive5.com/uparse/) and the chimeric sequences were identified and removed using UCHIME. The taxonomy of the 16S rRNA gene sequences was analyzed using the ribosomal database project (RDP) classifier algorithm (http://rdp.cme.msu.edu/) against the Silva (SSU123) 16S rRNA database using a confidence threshold of 70%.

#### Anammox bacteria purification analysis with fluorescence in situ hybridization

The purified anammox bacteria were fixed with ice in 4% paraformaldehyde for 12 h and were then dehydrated in 50%, 80%, and 100% ethanol for 10 min; 1*µ*L of probe Amx368 and 9 *μ*L of hybridization buffer were added to the enriched anammox bacteria in the hybridization wells(32, 33). The probe Amx368 CCTTTCGGGCATTGCGAA labeled with Cy3 at the 5’ end was applied to determine the total anammox bacteria. The hybridization was performed in a hybaid oven at 46 °C for 90 min. The total biomass of the enriched bacteria was stained with 10*μ*L DAPI at 5 *μ*g/ml for 15 min.(34) All operations related to the fluorochromes were conducted in a dark environment to avoid fluorescence quenching. The fluorescence images of the anammox bacteria and total biomass were obtained with a confocal laser scanning microscope (CLSM) (Olympus, FV1200); the excitation wavelength of Cy3 was set at 552 nm and the DAPI dye was set at 488 nm for the CLSM analysis. The ratio of the anammox to the total biomass was determined using Photoshop software.

### Image analysis of the anammox bacteria

#### Synchrotron soft X-ray 3D imaging of the anammox bacteria

The synchrotron soft X-ray imaging experiment was performed at the BL07W beamline at the National Synchrotron Radiation Laboratory (NSRL) in Hefei, China.(35) The purified anammox bacteria were centrifuged at 5000 g (Eppendorf, 5804R) for 2 min and were re-suspended in deionized water and then diluted to 1-2×10^7^/ml; 0.4 µL of the anammox bacteria suspension was seeded onto 100-mesh carbon-film copper grids. The grids with the anammox cells were plunged into a freezer for a few milliseconds to vitrify the anammox cell so that ice crystal contamination and structural damage of the anammox cells were avoided. Subsequently, the cells were rapidly placed into liquid nitrogen to remain vitrified to prevent radiation damage (36). The copper grids with the purified anammox cells were transferred into the vacuum cryogenic chamber of the soft X-ray instrument for imaging; 0.52 Kev photon energy was used to take advantage of the high natural contrast of the organics in the “water window” (37). A visible light microscope and a soft X-ray microscope were coupled to determine the suitable samples. The anammox cells perpendicular to the beamline (at 0° tilt) were the targets for the imaging. The images were acquired at a 2 s exposure time from -65°to 65° at 1°intervals, which was achieved by rotating the sample stage. The schematic of the 3D imaging of the intact anammox cell is shown in Fig 1. The synchrotron soft X-ray was focused on the condenser and interacted with the anammox bacteria sample on the sample stage. After adsorption and scattering by the anammox bacteria, the transmission of the soft X-ray signal was amplified by a condenser zone plate and detected by the charge coupled device (CCD).

**Fig 1.**
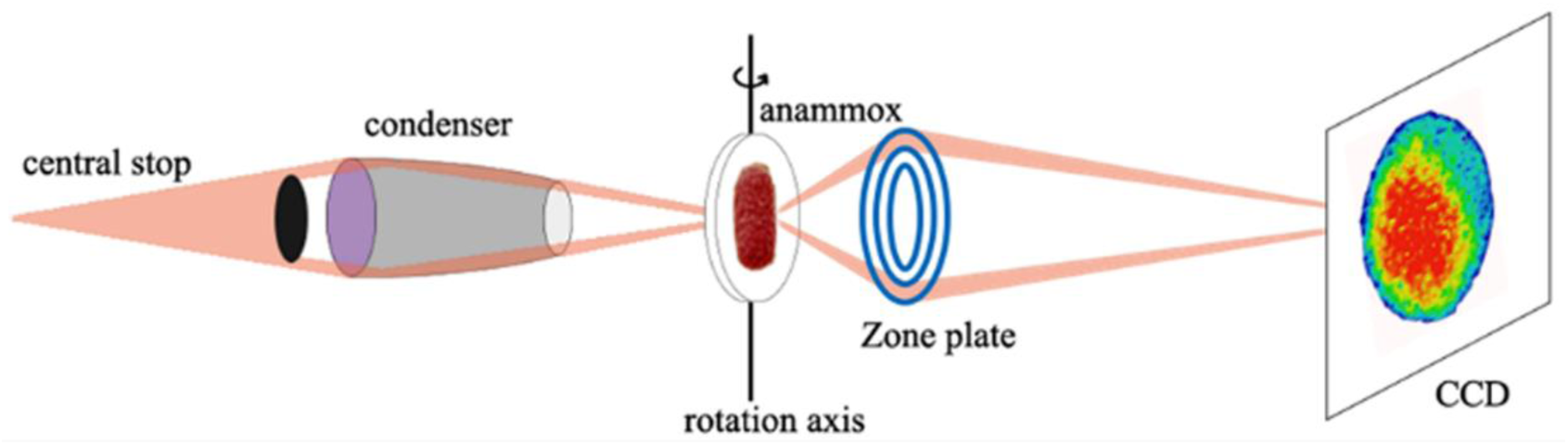
Schematic of the synchrotron soft X-ray nano-CT imaging process of the anammox cell. (The synchrotron soft X-ray was focused on the condenser, absorbed and scattered by the anammox bacteria, then amplified by a condenser zone plate and detected by the charge coupled device (CCD).

#### Reconstruction and segmentation of the anammox cells

To obtain the 3D information of the anammox ultrastructure, a tomographic reconstruction of the 131 images obtained from different angles was conducted using the FBP algorithm. Some angles were missing because of grid shading and the algebraic reconstruction technique (ART) was used for the missing wedges. The TV-SART in the TomoJ software was applied to obtain detailed structural information (38). The reconstruction results of the TV-SART algorithm were evaluated using the LAC values. The LAC represents the inherent ability of a substance for absorption the soft x-ray (39), and the images depicting the LAC were segmented to identify the different organelle structures and chemical composition (40, 41). Amira software was used to segment the reconstructed anammox bacteria cell, calculate the LAC values, measure the number of pixels of the anammoxosome organelle and anammox bacteria cell, and to create movies (26, 42).

#### Anammox imaging with TEM

The anammox bacteria were then fixed in 4% formaldehyde in a phosphate buffer solution on ice overnight. Subsequently, they were dehydrated in graded ethanol (50%, 70%, 80%, 90%, 95% and 100% ethanol). Next, the anammox bacteria were embedded in Epon resin and cut into 70nm thick slices using an ultramicrotome (Leica EM UC7)(10) and the sections were transferred to 100-mesh copper grids for the TEM analysis. The TEM imaging was performed with a HITACHI HT7700 instrument at an accelerating voltage of 100 kV.

### Morphological characteristics of the anammox bacteria response to Fe^2+^

Among the environmental conditions affecting anammox bacteria, iron has attracted the most attention. Iron plays a key role in anammox bacteria growth and affects the anammox activity and nitrogen removal capacity. To explore the shape adaptation and mechanism of the anammox bacteria response to different iron conditions, batch tests were performed using 120-ml serum bottles. 2g of the anaerobic granular sludge and 100 ml of the medium were added to the serum bottle. The medium content was the same as the EGSB medium except that the ammonium and nitrite were 100 mg/l and the concentrations of FeSO_4_ were 0, 0.015, 0.03, and 0.06 mM in different serum bottles, respectively. The medium in the serum bottle was aerated with argon for 10 min to ensure that the dissolved oxygen concentration was below 0.02 mg/l. The serum bottles were incubated in a shaking bath at 30 °C and 120 rpm. The medium in the serum bottle was changed every 24 hours and the batch tests lasted three months.

## Results

### Characteristics of anammox bacteria

The anammox granules sampled from the EGSB reactor are shown in Fig 2(a). The round granules were deep red and the diameter range was 1.2 mm to 6.0 mm. The microbial diversity analysis of the anammox granular sludge at the genus level is shown in Fig 2(b). The anammox bacteria included three genera: *Candidatus Jettenia, Candidatus Brocadia, and Candidatus Kuenenia,* accounting for 26.36%, 4.22% and 1.40% of the total microbial biomass in the EGSB reactor, respectively. The purified anammox cells had about 95% purity based on the Percoll density gradient centrifugation (Fig 2(c)). Although three genera of anammox bacteria were observed in the reactor, these three genera of *Candidatus Jettenia, Candidatus Brocadia, and Candidatus Kuenenia* had similar structural characteristics, including three separate compartments: periplasm, cytoplasm, and anammoxosome. Furthermore, the anammoxosome volume ratio in these three genera accounted for more than 60% of the anammox bacteria volume (11).

**Fig 2.**
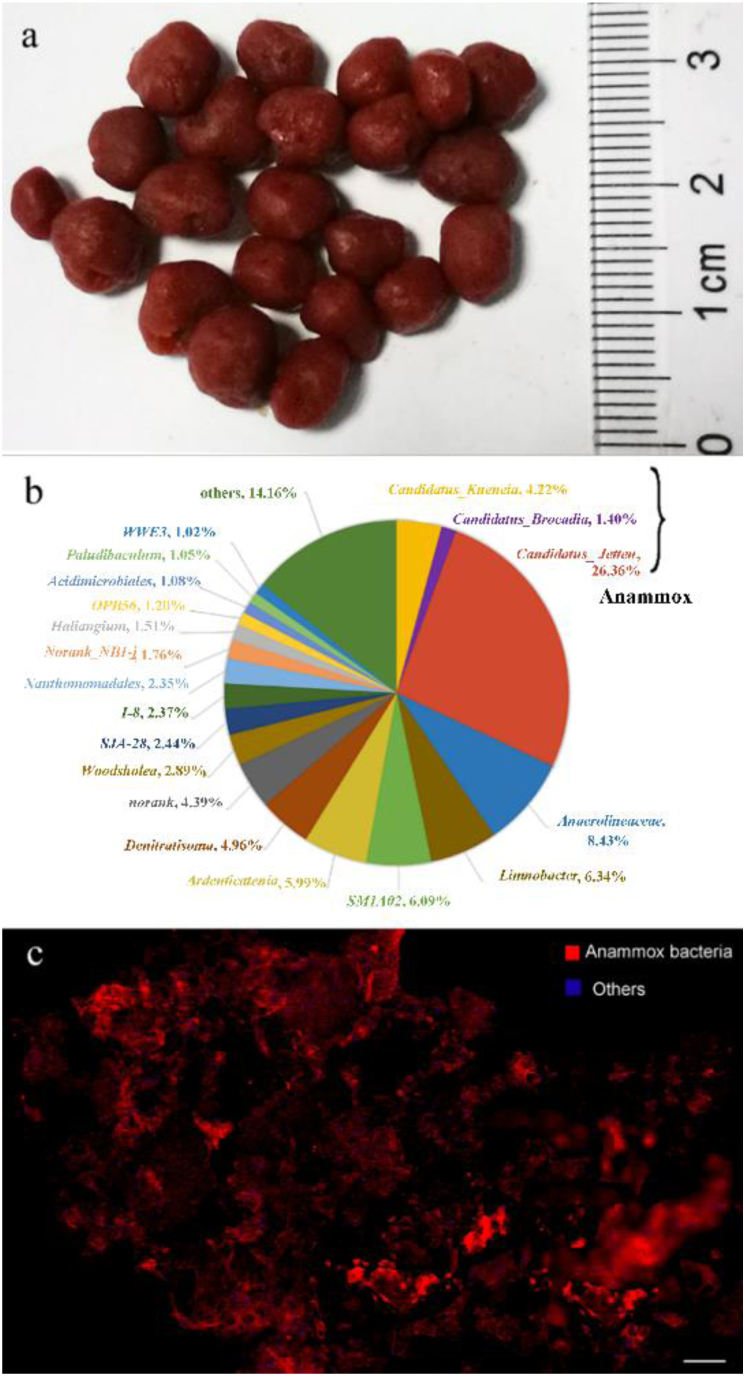
The anaerobic granular sludge in the EGSB reactor. (a) Anaerobic granular sludge sampled from the EGSB reactor. Scale bar shows 20 mm. (b) Microbial diversity analysis of the anaerobic granular sludge. (c) Confocal laser scanning microscope image of the anaerobic granular sludge. (Scale bar shows 200 *μ*m.) (The probe Amx368 targeting the anammox bacteria is shown in red and the DAPI dye targeting the total biomass minus the anammox bacteria is shown in blue.)

### Synchrotron soft X-ray transmission imaging and reconstruction of anammox bacteria

Fig S1 shows the synchrotron soft X-ray projection image of the intact cells; 131 projection images of the anammox bacteria at different angles were collected, and the 3D structure of the anammox bacteria was reconstructed using the FBP algorithm in the original soft X-ray microscopy system as shown in the Supplementary Movie 1. Fig S2 (a) shows a screenshot of the anammox bacteria. The structure inside the anammox bacteria was inferred to be anammoxosome. The slice images of the anammox bacteria are shown in Fig S2 (b) and the tomogram of the anammox bacteria is shown in Supplementary Movie 2. The soft X-ray absorption of all areas inside the anammox bacteria can be visualized and the grayscale difference is shown in the slice image.

To further investigate the interior ultrastructure of the anammox bacteria, the TV-SART (38) algorithm was used to reconstruct the anammox bacteria. The anammox bacteria in Fig S2 have a strong and heterogeneous high-absorption structure but the absorption difference inside the high-absorption membrane could not be distinguished due to limitations of the FBP algorithm. In addition, some artifacts were observed in the greyscale 3D reconstruction video (Supplementary Movie 1) and were attributed to missing angles. The TV-SART algorithm can compensate for these drawbacks and provides the LAC values.

The 3D reconstruction of the intact anammox bacteria using the TV-SART algorithm is shown in Supplementary Movie 3. Fig 3 shows the reconstructed anammox bacteria cell at different rotation angles with 60°intervals. The colormap ranges from 0 to 0.46 and represents the LAC value corresponding to the differences in the structure or composition. The LAC reconstruction provided a clearer representation of the ultrastructure of the anammox bacteria. The cluster (in yellow) inside the anammoxosome membrane had a higher LAC value and some nanoparticles (in green) had extremely high LAC values (Fig 3). Fig 4 shows the slice image of the anammox bacteria at different depths from 100 nm to 800 nm and the black arrows in Fig 4 indicate the nanoparticles with extremely high LAC values. The tomograph of the anammox bacteria reconstructed with the TV-SART algorithm is shown in Supplementary Movie 4. The slice image and video show the LAC values of all areas inside the anammox bacteria. It is noteworthy that some areas inside the anammoxosome membrane had low LAC values. In other words, the structure inside the anammoxosome membrane was relatively dense but overall, the areas of high soft X-ray absorption were discontinuous and inhomogeneous.

**Fig 3.**
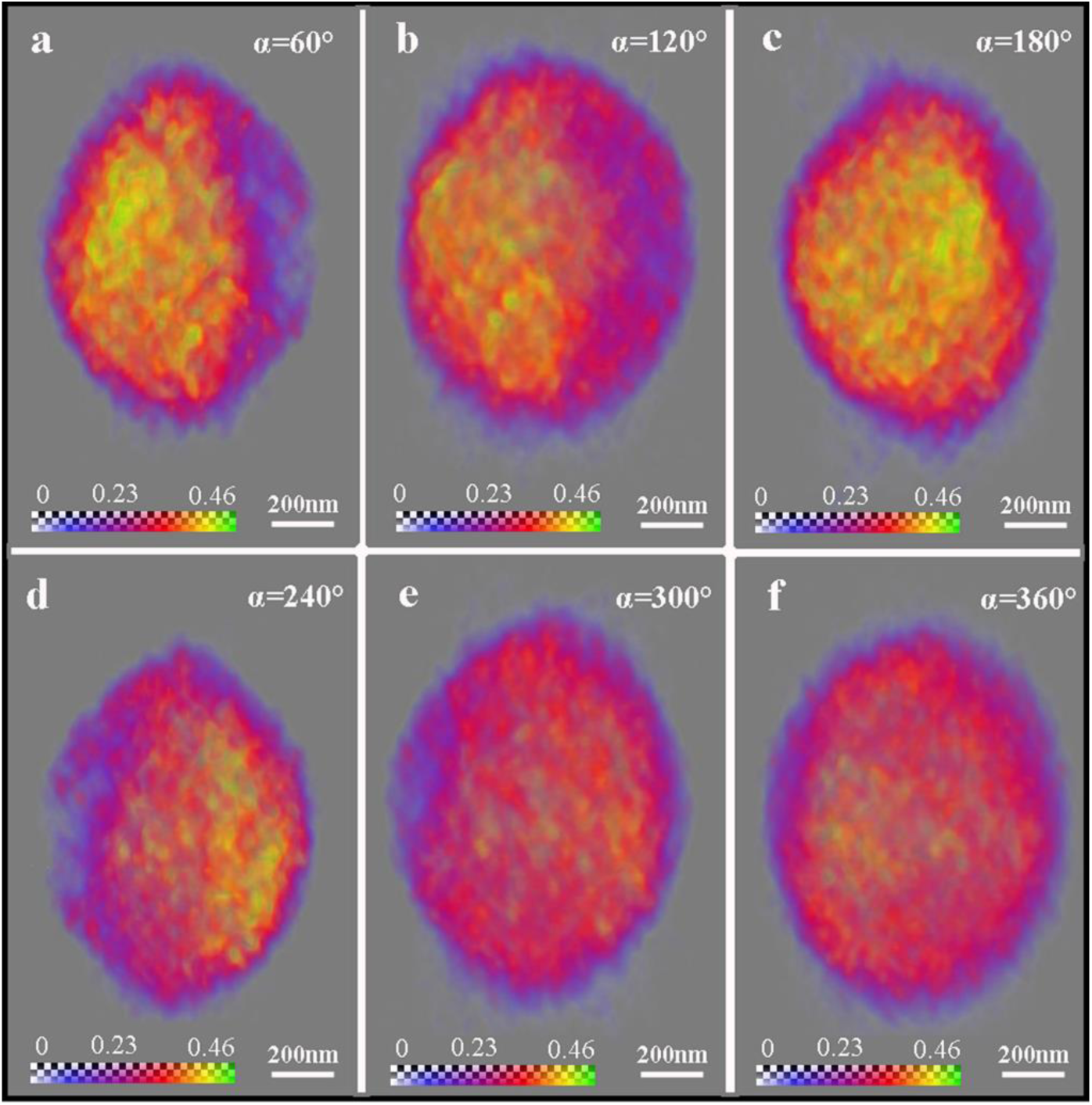
The intact anammox cell reconstructed using the TV-SART algorithm. (a), (b), (c), (d), (e), and (f) correspond to 60°, 120°, 180°, 240°, 300°, and 360°, respectively. (3D reconstruction of anammox bacteria at different rotation angles with 60°intervals.)

**Fig 4.**
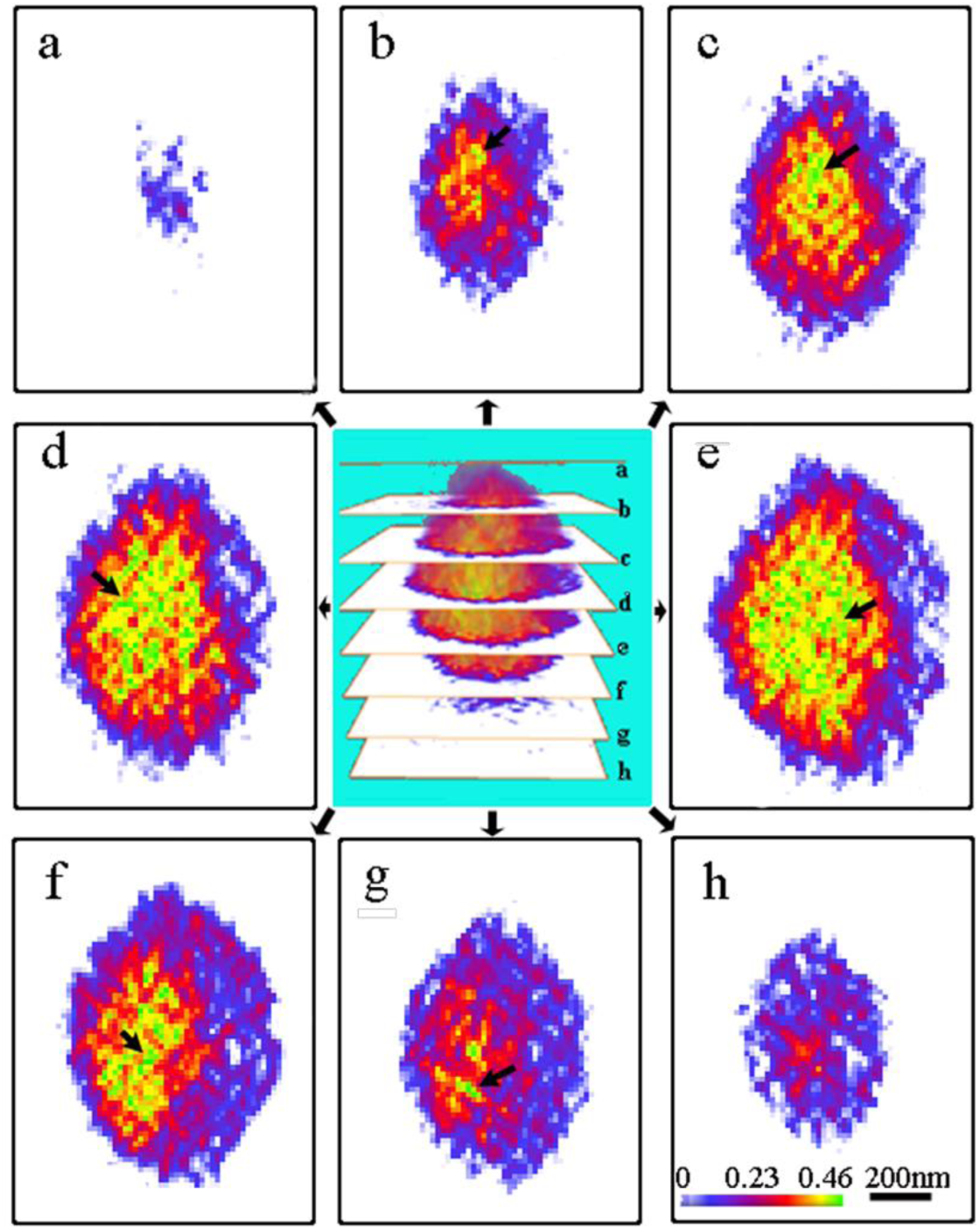
Slice images of the anammox bacteria cell at different depths. (a), (b), (c), (d), (e), (f), (g), and (h) correspond to slice depths at 100 nm, 200 nm, 300 nm, 400 nm, 500 nm, 600 nm, 700 nm, and 800 nm respectively (The black arrow points to nanoparticles in green. The scale bar and colormap in Figure h apply to all images).

### Quantitative and statistical analysis of the anammox bacteria

Since different biochemical components have specific LAC values, the organelles and other cell structures can be distinguished visually based on the differences in the biochemical composition and density (39, 42, 43). Thus, the image of the ultrastructure of the anammox cell was segmented based on the local LAC values using the Amira software, and the average LAC values of the anammox cell, the anammoxosome, and the nanoparticles were calculated to be 0.326±0.001 *μ*m^-1^, 0.389±0.001 *μ*m^-1^, and 0.460±0.001 *μ*m^-1^ respectively. The segmentation images of the anammox structure at different rotation angles are shown in Fig 5. The video of the segmentation of the 3D structure is shown in Supplementary Movie 5. The 3D segmentation structure of the anammox cell showed a distinct asymmetric structure, especially at the rotation angles of 60°and 240°(Fig 5(a) and Fig 5(d)), which indicated the importance of the 3D imaging of the intact anammox bacteria cells. Owing to the asymmetric distribution of the anammoxosome, the nano-scale slices observed by TEM at certain angles or depths do not represent the structure of the whole anammox bacteria, and different projection angles can cause changes in the anammoxosome volume ratio.

**Fig 5.**
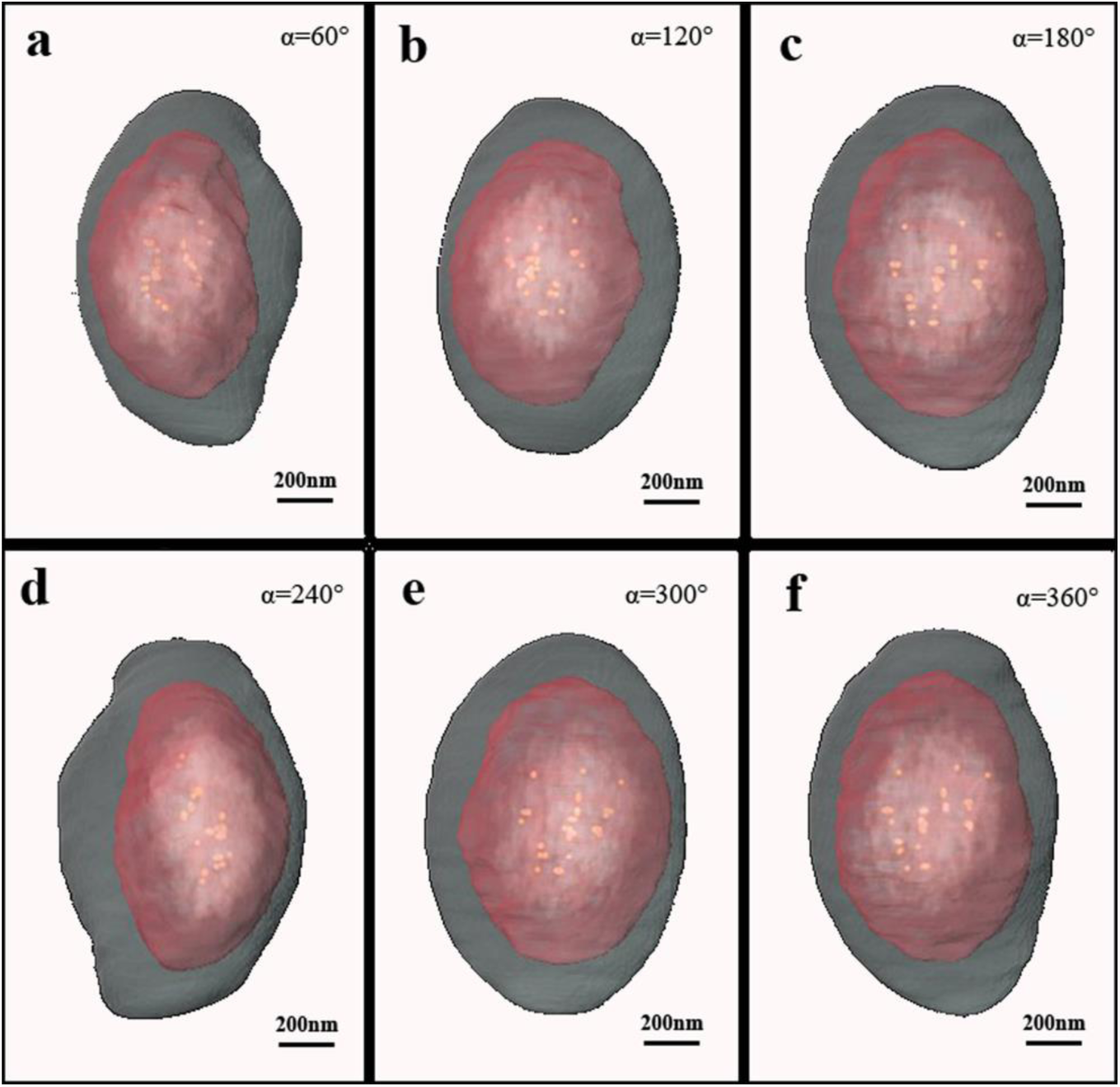
The segmentation images of the anammox cell at different rotation angles. (a), (b), (c), (d), (e), and (f) correspond to 60°, 120°, 180°, 240°, 300°, and 360°, respectively. (α represents the rotation angle).

To quantify the anammoxosome volume ratio, anammox cells were reconstructed and statistically analyzed using the Amira software. The ratio of the anammoxosome volume to the anammox bacteria cell volume (A/C) was 47±2.5% (Fig S3). This value was much lower than the previously reported volume ratio of 50%–80% with a large average error (11); the value was based on the area ratio of a nanoscale slice in a TEM ultrathin section (11). However, the values of A/C varied widely due to the exocentric anammoxosome structure. To verify the structure of the anammox bacteria, TEM images of the anammox bacteria slices were obtained in this study. Fig S4 (a), (b), and (c) represent different distributions and sizes of the anammoxosome. The slice in Fig S4 (b) indicates that the anammoxosome organelle account for the majority of the cell volume; however, the slice in Fig S4 (c) shows otherwise although this was not the most extreme case. In fact, some nanoscale slices show no anammoxosome. This may result in large differences in the A/C value. In this study, we calculated the volume ratio for the total number of pixels, i.e., the ratio of the volume of the anammoxosome pixels to the number of anammox cell pixels. The calculated volume ratio of the 3D reconstruction was almost equal to the actual value. To quantify the exact position of the anammoxosome in the anammox cell, we used the parameter of eccentricity (*e*), which is defined as the ratio of the longest distance to the shortest distance from the anammoxosome membrane to the cell membrane; this parameter reflects the degree of eccentricity of the anammoxosome. The maximum and minimum eccentricity values of the anammoxosome are shown in Fig S5. The average eccentricity is 3.25±0.43 (Fig S3); this number demonstrates the extent to which anammoxosome deviates from the center of the anammox bacteria. Although TEM imaging also provides information on the eccentricity of anammox bacteria (de Almeida et al. 2015), this method requires complex preprocessing, such as fixing, dehydrating, embedding, and sectioning, resulting in changes in the morphology of the anammox bacteria cell. Synchrotron soft X-ray imaging requires no pretreatment and the 3D nondestructive imaging of anammox bacteria cells can be performed in in-situ in the water. Thus, the results of the anammoxosome volume ratio obtained by synchrotron soft X-ray imaging is closer to reality than that obtained by TEM. Further, the degree of eccentricity of the intact anammoxosome can be calculated from the synchrotron soft X-ray images but has not been determined from TEM images.

### Morphological analysis of the anammox bacteria response to Fe^2+^

Iron is abundant in anammox bacteria and plays an important role in the anammox process in the form of free iron, iron-binding proteins, or iron-containing nanoparticles. The shapeshifting and mechanism of the response of the anammox bacteria to iron were investigated in this study. Fig 6(a) shows the average ratio of the anammoxosome volume to the anammox bacteria cell volume (A/C) at 0, 0.015, 0.03, and 0.06 mM FeSO_4_ respectively. It was evident that the A/C volume ratio decreased with increasing iron concentration from 0 to 0.06 mM. Accordingly, the LAC value changed with the A/C volume ratio (Fig 6(b)). In the absence of FeSO_4_ and in the presence of 0.015 mM FeSO_4_, the LAC value of the anammoxosome was 0.220 and 0.301 respectively, representing 43% and 23% decreases in the LAC value compared to the control group at 0.03 mM FeSO_4_. This may be related to two factors; one is the increase in the anammoxosome volume, which, in turn, leads to a decrease in the density of the anammoxosome. The other factor is that a low iron environment results in less iron accumulation in the anammoxosome, thus reducing the soft x-ray absorption ability. However, this is not applicable to the response at a concentration of 0.06 mM, which is likely due to the flexibility of the anammoxosome and the composition inside the anammoxosome outflow in response to the excessive iron concentration. This result provides a new perspective of the inhibitory effect of excessive iron concentrations on anammox bacteria. A similar response of anammox bacteria to iron has previously been observed using the TEM method (44). An increase in the iron concentration from 0.03 mM (Figure 6(c)) to 0.075 mM (Figure 6(d)) resulted in a decrease in the electron absorption density and area of the anammoxosome (the dark regions“D”in Figure 6 (d) and (d)).

**Fig 6.**
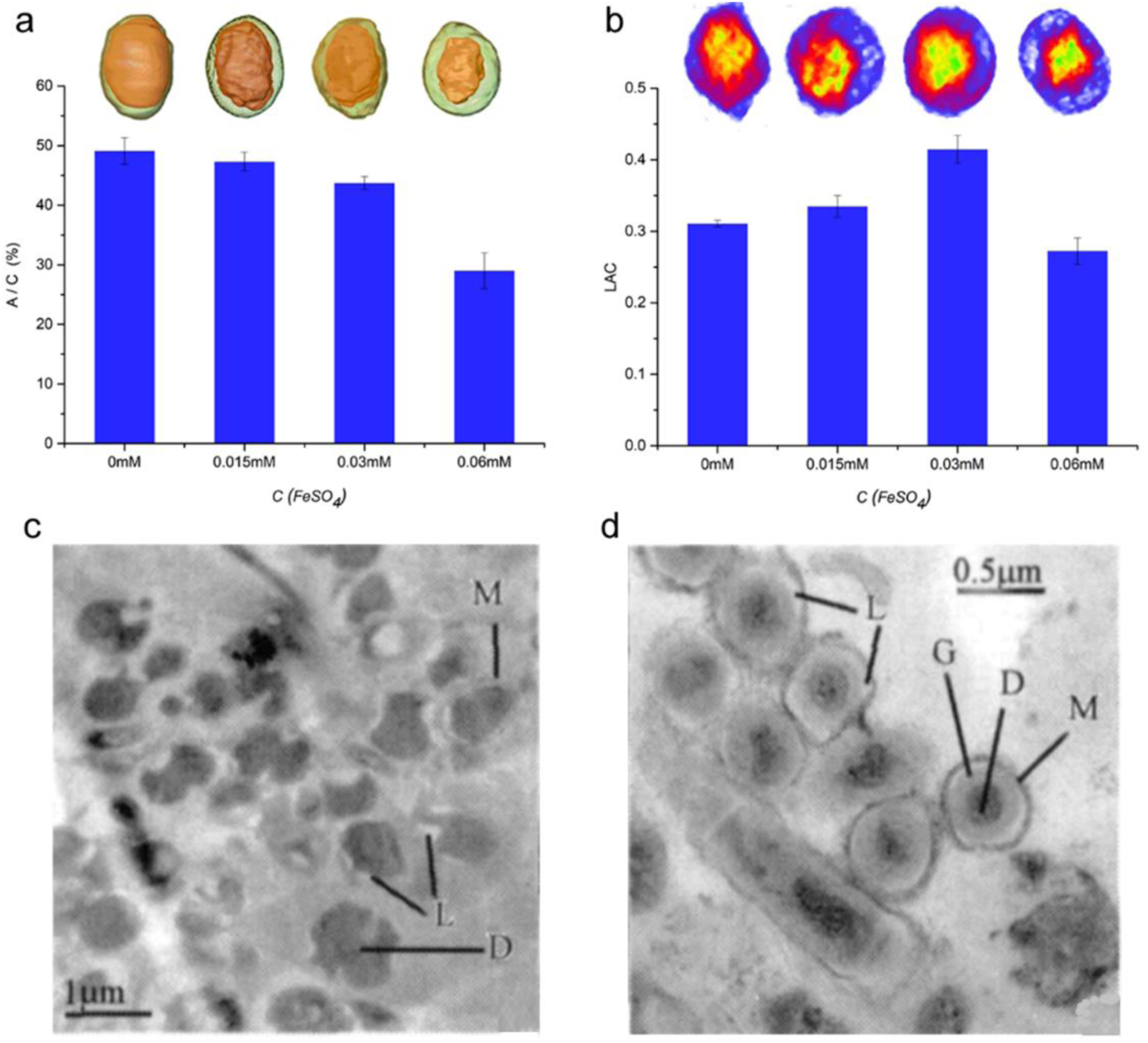
The effect of Fe^2+^ on the morphology of anammox bacteria. (a)The average A/C volume ratio at iron concentrations of 0, 0.015 mM, 0.03 mM, and 0.06 mM respectively. (b) LAC values at iron concentrations of 0, 0.015 mM, 0.03 mM, and 0.06 mM respectively. (c) and (d) are corresponding to the TEM imaging at the iron concentration of 0.03mM and 0.075mM (44). (Morphological analysis of anammox bacteria response to Fe^2+^)

## Discussion

Taken together, in this study, the synchrotron soft X-ray nano-CT imaging coupled with TV-SART reconstruction is used in the anammox bacteria morphology research for the first time. This novel method provided three-dimensional and interior ultrastructure information of the intact anammox bacteria in situ state.

The LAC value of the anammoxosome is much higher than that of the yeast organelles. The average LAC values of the chondriosome, vacuole, cell nucleus, and nucleoli were in the ranges of 0.34–0.38, 0.14–0.29, 0.25–0.27, and 0.32–0.34 *μ*m^-1^, respectively (42). The soft X-ray absorption of the specimen follows the Beer-Lambert Law. The LAC values represent the soft X-ray absorption intensity, and can be calculated using the following equation (1) (39):

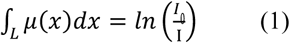

where *I*_0_ is the initial X-ray intensity and I is the output X-ray intensity. *μ*(x) is the local LAC value, which can be calculated by an iterative reconstruction using the TV-SART algorithm (40). The LAC value was determined based on the mass absorption coefficient and the density of the composition, as defined in the following equation (2):

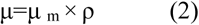

where *μ* is the LAC, *μ*_m_ is the mass absorption coefficient, and ρ is the density of the composition. Therefore, an area with a high LAC value is an area with a high mass absorption coefficient or high density(39, 43). The extremely high LAC value of the anammoxosome organelle indicated that the biochemical composition and structure of the anammoxosome were denser than that of the conventional organelles. Studies have shown that the anammox membrane contains abundant dense ladderane lipids with 1.5 g/cm^3^ density (14), resulting in higher LAC values of anammoxosome membrane than those of the cytoplasm. The density of the anammoxosome membrane is related to its functions, because the dense anammoxosome membrane plays an important role in preventing the leakage of valuable protons and intermediates during the anammox process. Furthermore, the anammoxosome is the site of the ammonia oxidation reaction that is associated with the presence of many enzymes. The high LAC values inside the anammoxosome membrane may be related to the numerous enzymes inside the anammoxosome, such as hydrazine dehydrogenase (HDH), hydrazine synthase (HZS), hydroxylamine/hydrazine oxidoreductase (HAO/HZO) (12), or metal elements (10).

Morphological analysis of the anammox bacteria response to Fe^2+^ provides a new perspective on underlying regulation mechanism of Fe^2+^ on anammox bacteria. The increased A/C ratio was the most characteristic shapes adaptation response of the anammox bacteria to Fe^2+^ hungry. An increase in the A/C ratio resulted in a larger surface area of the anammoxosome. Further, the larger surface area increased the number of metal binding sites (45). Thus, the shapeshifting of the anammox bacteria due to the increased A/C ratio at lower iron concentrations was presumed to enhance the iron absorption in the face of iron starvation. Iron is a critical and abundant element in anammox bacteria. The dark red color of the anammox bacteria is due to the abundance of iron and numerous iron-binding proteins (10, 46, 47). *Scalindua sp*. and *K. stuttgartiensis* can use iron and manganese oxides as electron acceptors and *K. stuttgartiensis* respires nitrate using iron as an electron donor (48, 49). In addition, these iron-binding cytochromes play an important role in electron transfer (46) and the oxidation-reduction process (47) in the energy metabolism of anammox bacteria. Finally, another interesting form of iron, i.e., iron-containing electron-dense nanoparticles was reported in the anammoxosome (10, 47). These nanoparticles are relatively abundant in iron and phosphorus (10). Thus, it is probable that the anammox bacteria adapt to an iron-limited environment by increasing the anammoxosome volume ratio to be able to absorb more iron. This shapes adaptation mechanism is similar to that of the Mn-oxidizing bacteria *Halomonas meridiana and Marinobacter algicola,* which adapt to Mn(II))-induced stress by increasing the cell length and volume to achieve better Mn (II) oxidizing ability (21). This shapeshifting phenomenon indicates a sensitivity and dependence of anammox bacteria to iron. In future studies, we will investigate this mechanism of iron regulation in the anammox process in more detail.

## Conclusion

Synchrotron soft X-ray nano-CT imaging coupled with TV-SART algorithm resulted in a breakthrough in intact anammox bacteria morphology research. This novel method provided high-contrast and high-resolution 3D images of the whole anammox bacteria cell in its natural state. The ultrastructure of the anammox bacteria was imaged and the linear absorption coefficient, anammoxosome volume ratio, and eccentricity were quantified. On this basis, the shape adaptation and mechanism of the anammox bacteria responses to Fe^2+^ were explored. Synchrotron soft X-ray nano-CT imaging technology provides a new perspective for the study of the biophysical properties of cells in different environments and engineering processes.

## Author contributions

All authors assisted with data interpretation and manuscript review. Y.-P.C, G.L, M-W.P, and Y. G conceived and designed the research. Y.-P.C and M-W.P analyzed the data. J.-H.L and L.C performed the TV-SART 3D reconstruction. H. W cultivated all the anaerobic granular sludge. H.-Y.L and P. L assisted with the synchrotron soft X-ray imaging experiments. Z.-Z. X conducted the TEM imaging experiments. P. Y purified the anammox bacteria and Y. S isolated the anammoxosome. Y.-P.C and G.L performed the segmentation of the cell based on the LAC value. J.-S. G designed all figures with F.F and with the help from all other authors. M.-W.P wrote the manuscript with help from all authors.

## ACKNOWLEDGMENTS

This work was financially supported by the National Natural Science Foundation of China (21876016 and 51578527), the Chongqing Science and Technology Commission (cstc2018jcyjAX0366), the Fundamental Research Funds for the Central Universities (2018CDQYCH0028), and the National Key Research & Development Program of China (2016YFE0205600). We gratefully acknowledge Bing-Hong Wan for the guidance on the Amira software, and the BL07W beamline at the National Synchrotron Radiation Laboratory in Hefei, China; the supports of Majorbio, China.

